## Supplementary Materials for "Clustering huge protein sequence sets in linear time"

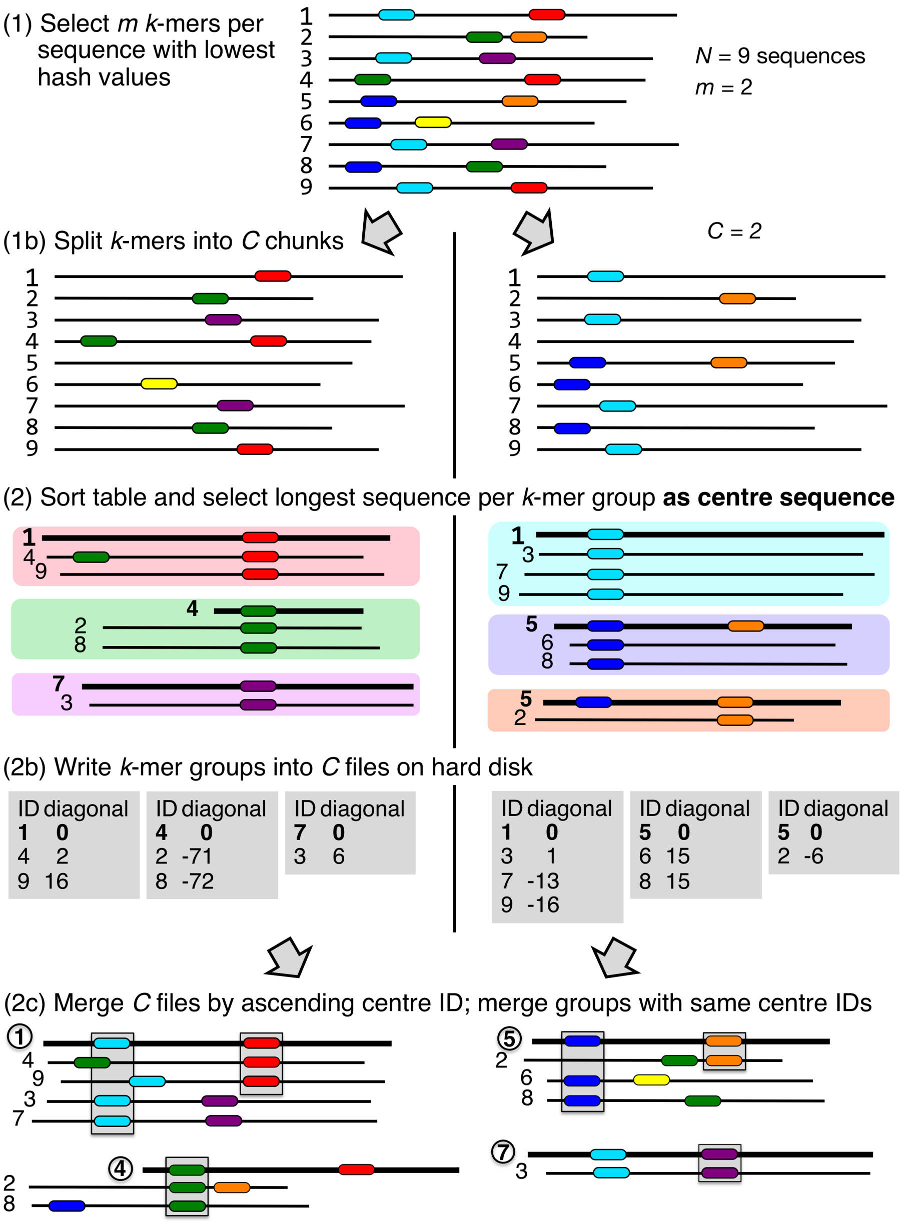


**Figure S** 1. **Splitting the database into chunks** For each chunk index $c \in \{0, \ldots, C-1\}$ we run Linclust steps 1 and 2 (Figure 1) normally but extract only $k$-mers whose numerical index modulo $C$ yields a rest $c$. This way each of the $C$ runs builds up a $k$-mer table with only about $mN/C$ lines instead of $mN$, and hence each run needs $C$ times less memory. Each run writes out a file with all found $k$-mer groups (sequence ID, diagonal). The diagonal is $i - j$ of the k-mer match, where $i$ is the position of the groups $k$-mer in the centre sequence and $j$ the position in the other sequence. Afterwards all $C$ files are merged into a single file such that $k$-mer groups are sorted by ascending centre IDs. Finally, Linclust steps 3 to 5 are performed as usual.

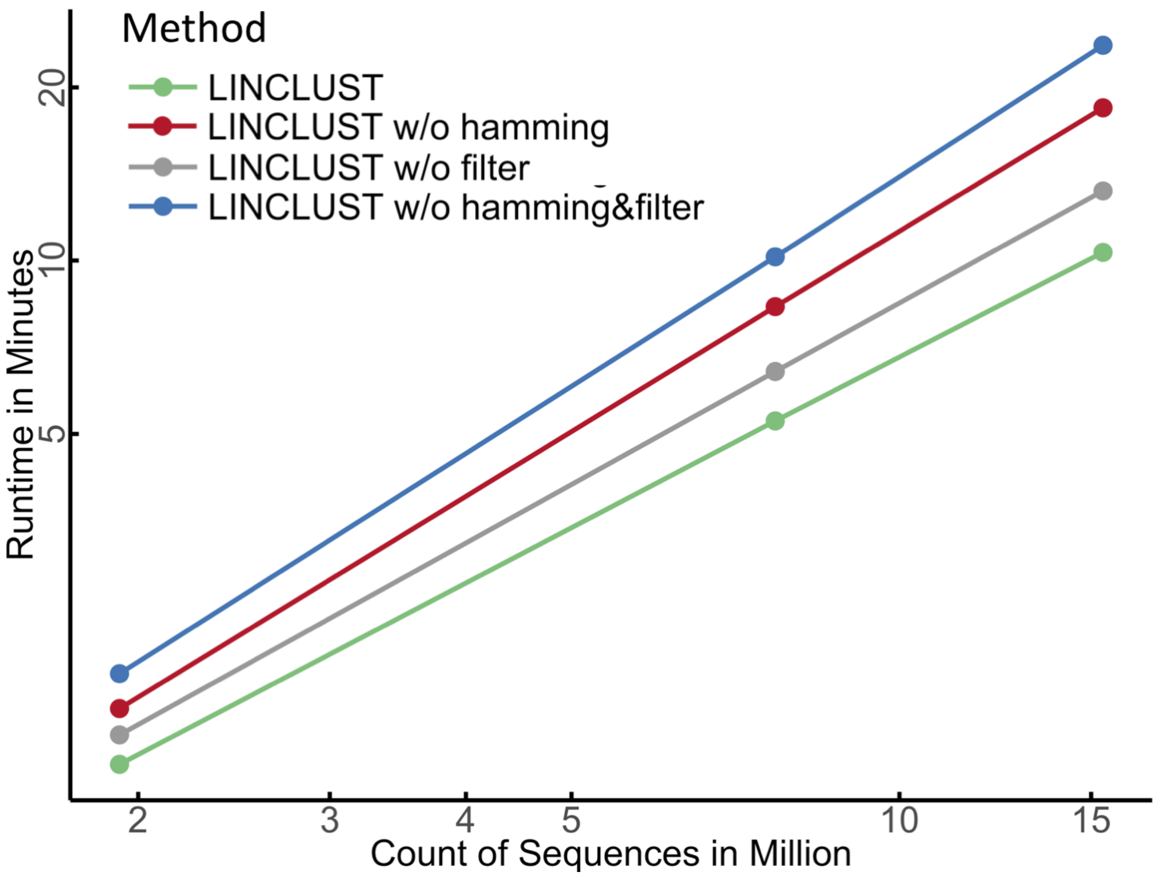


**Figure S** 2. **Contribution of Hamming distance pre-clustering and gapless local alignment filter steps to the decrease of the Linclust runtime** Double-logarithmic plot of runtimes versus sequence set size illustrating the contribution of runtime decrease for the Hamming distance pre-clustering and ungapped alignment filtering stage.

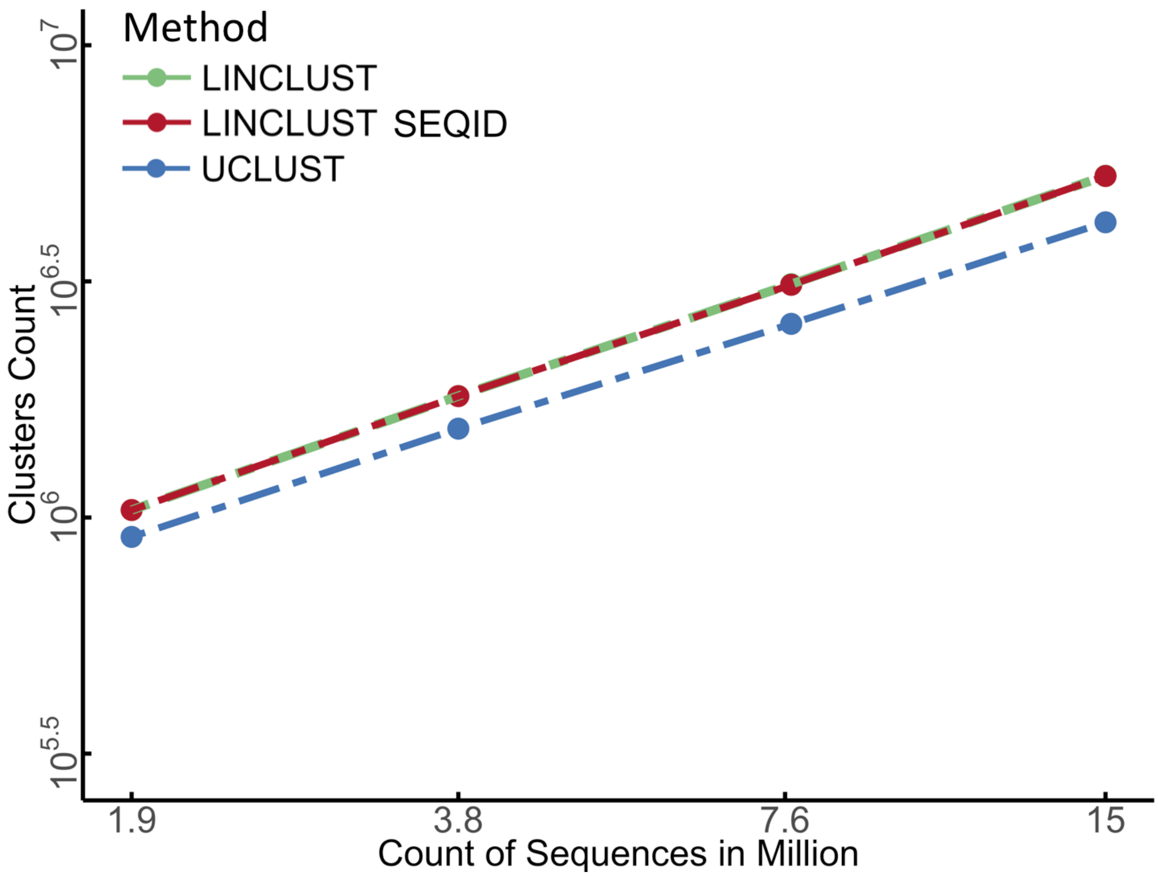


**Figure S** 3. **Equivalence of two sequence identity measures.** Number of clusters obtained at 50% sequence identity for Linclust with two different sequence identity definitions: (1) the default definition ("Linclust"), based on the local alignment score divided by the maximum length of the two aligned sequence segments; and (2) the fraction of identical residues in the alignment relative to the number of aligned columns including gaps ("Linclust seqid"). Both measures produce very similar numbers of clusters.

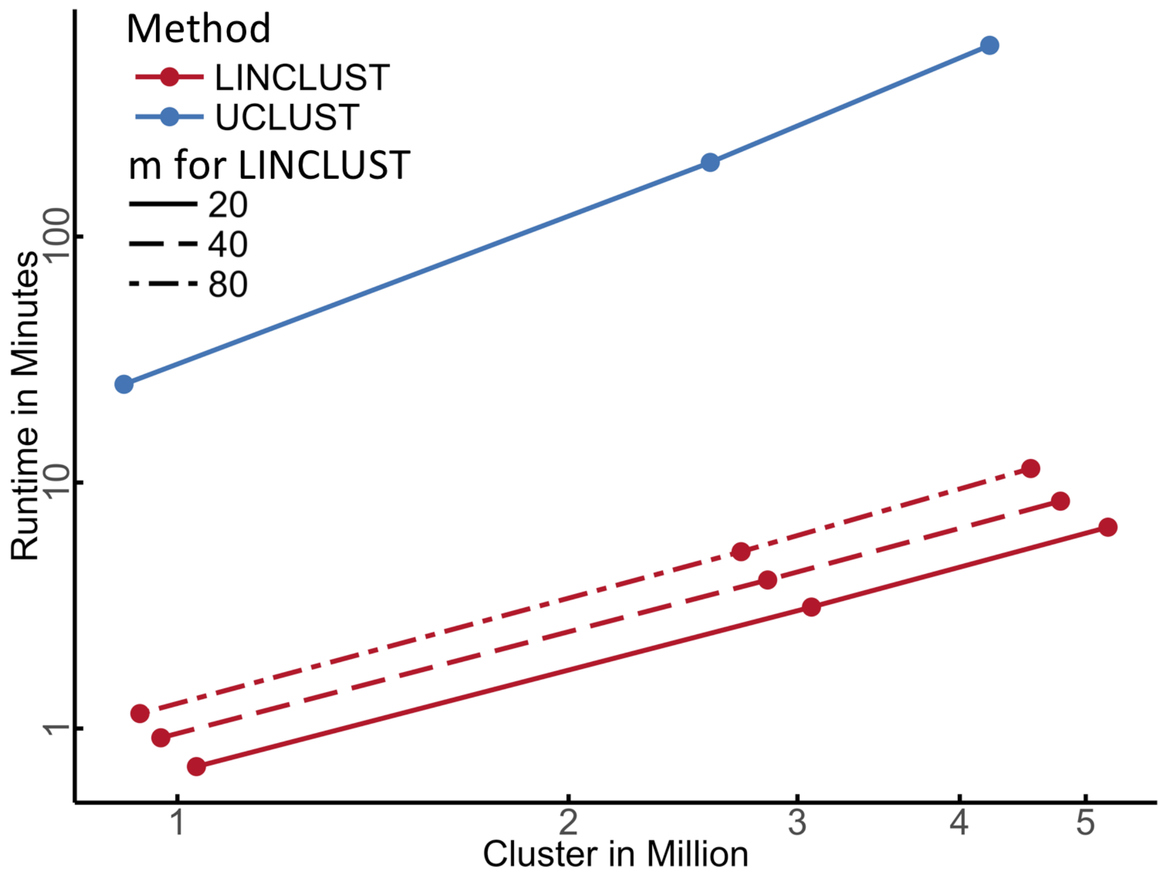


**Figure S** 4. **Influence of the number $m$ of $k$-mers extracted per sequence.** Double-logarithmic plot of runtimes versus cluster size. Through the parameter $m$, the number of $k$-mers selected per sequence, the user can set the trade-off between sensitivity and speed. At $m = 80$ the runtime of Linclust increases by a factor of 1.66 over the default setting $m = 20$ while producing 8% fewer clusters. At $m = 80$ , Linclust generates only 7% more clusters than UCLUST while still being much faster.

```c
// (c) 2017 Johannes Soeding & Martin Steinegger, Gnu Public License version 3
// Rotate left macro: left circular shift by numbits within 16 bits

#define RoL(val, numbits) (val << numbits) ^ (val >> (16 - numbits))

// Transform each letter x[i] to a fixed random number RAND[x[i]]
// to ensure instantaneous mixing into the 16 bits
// Do XOR with RAND[x[i]] and 5-bit rotate left for each i from 1 to k
unsigned circ_hash(const int * x, unsigned length){
 short unsigned RAND[21] = {0x4567, 0x23c6, 0x9869, 0x4873, 0xdc51, 0x5cff, 0x944a, 0x58ec,
                            0x1f29, 0x7ccd, 0x58ba, 0xd7ab, 0x41f2, 0x1efb, 0xa9e3, 0xe146,
                            0x007c, 0x62c2, 0x0854, 0x27f8, 0x231b}; // 16 bit random numbers
 short unsigned h = 0x0;
 h = h^ RAND[x[0]];                  // XOR h and ki
 for (int i = 1; i < length; ++i){
   h = RoL(h, 5);
   h ^= RAND[x[i]];                  // XOR h and ki
 }
 return h;
}

// Rolling hash variant for previous hash function:
// Computes hash value for next key x[0:length-1] from previous hash value
// hash( x[-1:length-2] ) and x_first = x[-1]
unsigned circ_hash_next(const int * x, unsigned length, int x_first, short unsigned h){
 short unsigned RAND[21] = {0x4567, 0x23c6, 0x9869, 0x4873, 0xdc51, 0x5cff, 0x944a, 0x58ec,
                            0x1f29, 0x7ccd, 0x58ba, 0xd7ab, 0x41f2, 0x1efb, 0xa9e3, 0xe146,
                            0x007c, 0x62c2, 0x0854, 0x27f8, 0x231b}; // 16 bit random numbers
 // undo INITIAL_VALUE and first letter x[0] of old key
 h ^= RoL(RAND[x_first], (5*(length-1)) % 16);
 // circularly permute all letters x[1:length-1] to 5 positions to left
 h =  RoL(h, 5);
 // add new, last letter of new key x[1:length]
 h ^= RAND[x[length-1]];
 return h;
}
```

**Figure S** 5. ***k*-mer hashing function implemented in C.**
